# Chemogenetic inhibition of the amygdala modulates emotional behavior expression in infant rhesus monkeys

**DOI:** 10.1101/534214

**Authors:** Jessica Raper, Lauren Murphy, Rebecca Richardson, Zoe Romm, Zsofia Kovacs-Balint, Christa Payne, Adriana Galvan

**Author notes:** **Corresponding Author**: Dr. Jessica Raper, Yerkes National Primate Research Center, 954 Gatewood Rd NE, Atlanta, GA 30329. **Author Contributions**: JR, LM, and CP Designed Research; JR, LM, RR, ZR, and AG Performed Research; JR, AG, and ZK-B Analyzed Data; JR, LM, CP, and AG Wrote Paper.

## Abstract

Manipulation of neuronal activity during the early postnatal period in monkeys has been largely limited to permanent lesion studies, which can be impacted by developmental plasticity leading to reorganization and compensation from other brain structures that can interfere with the interpretations of results. Chemogenetic tools, such as DREADDs (designer receptors exclusively activated by designer drugs), can transiently and reversibly activate or inactivate brain structures, avoiding the pitfalls of permanent lesions to better address important developmental neuroscience questions. We demonstrate that inhibitory DREADDs in the amygdala can be used to manipulate socioemotional behavior in infant monkeys. Two infant rhesus monkeys (1 male, 1 female) received AAV5-hSyn-HA-hM4Di-IRES-mCitrine injections bilaterally in the amygdala at 9 months of age. DREADD activation after systemic administration of either clozapine-N-oxide or low dose clozapine resulted in decreased freezing and anxiety on the human intruder paradigm and changed the looking patterns on a socioemotional attention eye tracking task, compared to vehicle administration. The DREADD-induced behaviors were reminiscent of, but not identical to, those seen after permanent amygdala lesions in infant monkeys, such that neonatal lesions produce a more extensive array of behavioral changes in response to the human intruder task that were not seen with DREADD-evoked inhibition of this region. Our results may help support the notion that the more extensive behavior changes seen after early lesions are manifested from brain reorganization that occur after permanent damage. The current study provides a proof-of-principle that DREADDs can be used in young infant monkeys to transiently and reversibly manipulate behavior.

**Statement of Significance:** Many neurodevelopmental disorders exhibit abnormal structural or functional amygdala development and alterations in socioemotional behavior. To date, developmental neuroscience studies have relied on permanent lesions techniques to investigate how atypical amygdala development impacts socioemotional behaviors, which may not adequately recapitulate the role of amygdala dysfunction in the manifestation of aberrant behavior. The present study sought to demonstrate that the designer receptors exclusively activated by designer drugs (DREADDs) chemogenetic tool could transiently inhibit amygdala activity in infant monkeys resulting in alterations in socioemotional behavior. This proof-of-principle study supports the use of chemogenetics for developmental neuroscience research, providing an opportunity to broaden our understanding of how changes in neuronal activity across early postnatal development influences behavior and clinical symptoms.

## Introduction

The amygdala plays an important role in social behavior and emotional responses to threats in the environment (Aggleton, 2000). This knowledge has largely come from research on adult humans and animals. Yet, the amygdala has a protracted postnatal development (Payne et al., 2010; Chareyron et al., 2012; Hunsaker et al., 2014), in which growth of the amygdala appears to coincide with refinement of social and threat detecting skills (Mendelson, 1982; Mendelson et al., 1982.; Kalin et al., 1991). Investigations of how amygdala development contributes to these skills have largely utilized permanent lesion techniques, finding that damage to the amygdala during the early postnatal period contributes to life-long changes in social, emotional, and neuroendocrine function (Bauman et al., 2004; Bliss-Moreau et al., 2010; Bliss-Moreau et al., 2011a; Bliss-Moreau et al., 2011b; Raper et al., 2013b; Raper et al., 2014b; Raper et al., 2014a; Stephens et al., 2015; Bliss-Moreau et al., 2017; Moadab et al., 2017). Given the potential for neural plasticity during this postnatal period, there is considerable opportunity for reorganization and compensation from other brain structures. In fact, several studies have demonstrated how lesions early in life contribute to such reorganization (Machado et al., 2008; Raper et al., 2014b; Grayson et al., 2017; Payne et al., 2017). In order to avoid the pitfalls of permanent lesions and better address these questions we need tools that transiently activate or inactivate the amygdala during early development.

Pharmacological inactivation techniques have been used to manipulate amygdala activity and influence behavior (Wellman et al., 2005; Wellman et al., 2016). However, these techniques require the surgical placement of skull-mounted chambers for intracerebral drug infusions, which are not ideal for developmental research on infant monkeys with rapidly growing brains and bodies. An alternative approach that would not require permanently mounted hardware nor rely on repeated intracerebral injections is greatly needed for developmental research.

Chemogenetic techniques provide powerful tools for behavioral neuroscience because they allow remote manipulation of neuronal activity (Sternson and Roth, 2014). Designer receptors exclusively activated by designer drugs (DREADDs) are the most commonly used chemogenetic tool. DREADDs involves the use of artificial G protein-coupled receptors that are synthetic variants of muscarinic receptors (Armbruster et al., 2007; Farrell and Roth, 2013; Roth, 2016), are not activated by acetylcholine (the endogenous ligand of muscarinic receptors) and are not constitutively active. Instead, DREADDs are exclusively activated with high efficacy by exogenous ligands. The available DREADDs include receptors that are coupled to Gi/o, Gq/11 and Gs proteins termed M4Di, M3Dq and GsD, respectively (Wess et al., 2013; Roth, 2016), providing tools that can either increase or decrease the activity of transduced neurons, depending on the intracellular pathway activated. The expression of DREADDs can be targeted to specific neurons using viral transfection (Urban and Roth, 2015; Roth, 2016). The most commonly used ligand to activate DREADDs is clozapine-N-oxide (CNO), although recent reports have shown that CNO can be reverse metabolized into clozapine (Raper et al., 2017; Allen et al., 2018). Thus, an alternative method to active the DREADDs is to use clozapine itself, at low doses that will only preferentially activate the DREADDs (Gomez et al., 2017).

DREADDs have been widely used in rodent models, but reports in nonhuman primates (NHPs) have been limited. To date, only four publications have reported DREADDs manipulation of behavior or neuronal activity in NHPs (Eldridge et al., 2016; Grayson et al., 2016; Nagai et al., 2016; Upright et al., 2018). While these studies provide critical information demonstrating the ability to use chemogenetic tools in NHPs to manipulate neuronal activity and impact behavior, they have all focused on adult monkeys. Considering its advantages over permanent lesions and other transient inactivation techniques (i.e., reversibility and minimal invasiveness), chemogenetics hold their greatest promise for developmental neuroscience research. The current study provides proof-of-principle that chemogenetic tools can be used to manipulate amygdala activity in infant monkeys.

## Materials and Methods

### Subjects

Two infant Indian rhesus macaques (*Macaca mulatta*), one male and one female (1.85 and 2.0 kg BW, respectively), were used in this study. The animals were raised by their mothers in large social groups at the Yerkes National Primate Research Center (YNPRC) Field Station (Lawrenceville, GA) until 7 months of age, when they were transferred to indoor pair-housing at the YNPRC Main Station (Atlanta, GA) for the duration of the study. The Animal Care and Use Committee of Emory University approved all procedures, which were performed in accordance with the National Institutes of Health Guide for the Care and Use of Laboratory Animals. Figure 1 outlines the study design, behavioral testing, and timing of ligand administrations.

**Figure 1.**
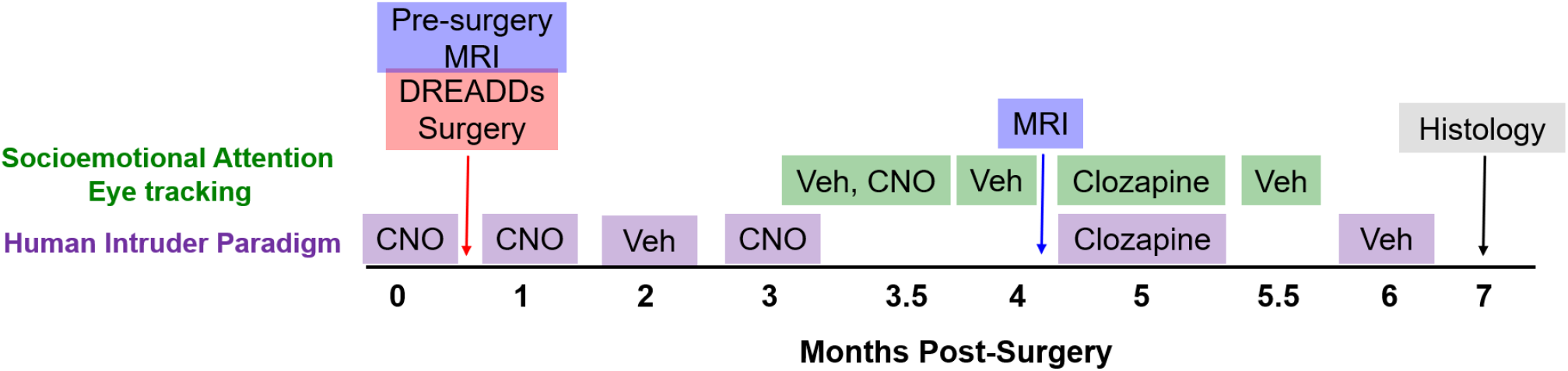
Schematic of the study design. Two infant rhesus monkeys received bilateral injections of AAV-hM4Di DREADD into the amygdala at 9 months of age, followed by 6 months of testing with either Vehicle (Veh), clozapine-n-oxide (CNO, 10mg/kg), or clozapine (0.1mg/kg). Prior to DREADD transduction, infants were tested on the Human Intruder paradigm with CNO to rule out any impact of the drug on the behavior in naïve animals. Magnetic resonance imaging (MRI) was conducted immediately prior to surgery to aid with amygdala injection localization and again at 4 months post-surgery in relation to another study not reported here. Monkeys were sacrificed at 7 months post-surgery to determine DREADD expression with histology.

Screening for AAV Neutralizing antibodies: Nonhuman primates’ native immunity has been shown to impact transduction in other CNS tissues (i.e., eye (Kotterman et al., 2015)) and neutralizing antibody response to multiple virus solution intracerebral injections have been directly linked to weak transduction in neurons (Mendoza et al., 2017). However, it is currently unknown whether native immunity to AAVs can impact transduction of neuronal tissue. Out of an abundance of caution, infant monkeys were screened for neutralizing antibodies to AAV5 prior to surgery to reduce the possibility that such antibodies might interfere with viral transduction.

Blood samples (2 ml) were collected in tubes containing EDTA (3.5 mg), centrifuged at 2500 rcf for 15 min, and plasma was pipetted off for assay by the Virology Core at the YNPRC (Atlanta, GA). AAV5 neutralization titers were determined by 50% infection inhibition assay (IC50) on HeLa cells (ATCC, Manassas, VA) freshly infected with Adenovirus serotype 5 (Ad5, ATCC, Manassas, VA). Briefly, plasma samples were heat inactivated for 1 hour at 56°C and diluted to 1:5, 1:20, 1:80, and 1:240. Plasma dilutions were then mixed with AAV5 bearing a luciferase expression construct (AAV5-luc, University of North Carolina Vector Core, Chapel Hill, NC) and incubated for 1 hour at 37°C. After initial incubation, plasma/AAV5-luc mixtures were added to the Ad5-infected HeLa cells and incubated at 37°C in 5% CO_2_ incubator for 48 hours. Luciferase expression was determined using the Bright-Glo Luciferase Assay system (Promega, Madison, WI) according to the manufacturer’s protocol, and measured on a BioTek Gen5 luminometer. IC50 was calculated relative to negative serum controls. Results revealed that neutralizing antibody titers for AAV5 were below the level of detection in the male and 1:20 for the female.

### Surgery

At 9 months of age, animals underwent DREADDs transduction surgery. Injection of inhibitory DREADDs bilaterally into the amygdala followed procedures similar to those described in previous studies of amygdala lesions in monkeys (Raper et al., 2014b). Briefly, on the day of surgery, the infants were removed from their cagemate, sedated with ketamine hydrochloride (1 mg/kg) and maintained with isoflurane (1–2% to effect). Their head was secured in a nonferromagnetic stereotaxic apparatus, and T1-weighted MRI sequences were acquired using a Siemens 3.0T/90 cm whole-body scanner and a 3 inch circular surface coil. A high resolution T1-weighted scan [spin-echo sequence, echo time (TE) = 11 ms, repetition time (TR) = 450 ms, contiguous 1 mm section, 12 cm field-of-view (FOV), 256 × 256 matrix] obtained in the coronal plane was used to determine the coordinates of injection sites in the amygdala. An additional T1-weighted MRI was acquired at 4 months post-surgery in the context of other studies (not reported here), following these same procedures.

All surgical procedures were performed under aseptic conditions, an intravenous drip (0.45% dextrose/0.9% NaCl) was placed to maintain normal hydration, and vital signs (heart rate, respirations, blood pressure, expired CO2) were monitored throughout the surgery. Nolvasan solution was used to disinfect the scalp and a local anesthetic (Bupivacaine 0.25% concentration, 1.5 ml) was injected subcutaneously along the midline to reduce the pain during skin incision. After the skin and underlying connective tissue were gently displaced laterally, two small craniotomies were made in front of bregma and above the amygdalae, and the dura was cut and retracted to expose the brain. Animals received viral vector injections of Dr. Bryan Roth’s DREADD construct AAV5-hSyn-HA-hM4Di-IRES-mCitrine (6 × 10^12^ vg/ml; Duke Viral Vector Core, Duke University, Durham, NC) administered in eight sites within the center of each amygdala using 10 μl Hamilton syringe. Needles were lowered simultaneously in both hemispheres, 5 μl were manually injected (0.4 μl/min) at each site for a total of 40μl per hemisphere. After each injection, a 3 min waiting period was allotted to minimize viral spread during needle retractions.

At the completion of the surgical procedures, the dura was closed with silk sutures, the bone opening was covered with Surgicel NU-KNIT (absorbable hemostat), and connective tissues and skin were closed. The animal was removed from anesthesia and placed in a temperature controlled incubator ventilated with oxygen until full recovery from anesthesia. All animals received banamine (1 mg/kg for 3 d), dexamethasone (0.5 mg/kg for 3 d), and antibiotic (rocephin, 25 mg/kg for 7 d) after surgery to prevent pain, edema, and infection, respectively. Behavioral testing began 5 weeks after surgery to allow animals to recover and provide adequate time for DREADD expression.

### Administration of Ligands and Plasma Assays

All ligands were administered through subcutaneous injection in the interstitial space between the shoulder blades of the animals. CNO was administered at 10mg/kg body weight, whereas clozapine was administered at 0.1μg/kg body weight. CNO powder (NIMH C-929, RITI International) was stored at −20 °C, protected from light. CNO solutions were freshly prepared on each experimental day, by dissolving the drug in 100% dimethyl sulfoxide (DMSO, Sigma-Aldrich, St. Louis MO, USA) and then diluting with calcium and magnesium-free phosphate-buffered saline (PBS, Corning, Corning, NY, USA) to a final concentration of 10 mg/mL in 15% DMSO. To prevent degradation due to light exposure, vials and syringes that contained CNO solution were wrapped in aluminum foil. Clozapine powder (Tocris, Minneapolis, MN) was stored at room temperature, protected from light. Clozapine solutions were also freshly prepared on each day of experiment, using the same dilution methods described for CNO. Percentages of DMSO and PBS remained constant across all injections to avoid any potential confound that might arise from differences in DMSO concentrations per injection. Vehicle injections consisted of 85% PBS and 15% DMSO.

Animals were trained for awake blood collection from the saphenous vein, following established protocols at the YNPRC (Raper et al., 2014b). On behavioral testing days, blood samples were collected several times after ligand administration to examine plasma concentrations of the ligand during behavioral tasks. For CNO administration, blood was collected at 20, 60, 90 and 135min post-injection. For clozapine administration, blood was collected at 2, 40, and 60 minutes post-injection. All blood samples (1-2ml) were collected in prechilled 2 ml tubes containing EDTA (3.5 mg) and immediately placed on ice. Samples were centrifuged at 2500 rcf for 15 min in a refrigerated centrifuge (at 4 °C). Plasma was pipetted off and stored at −80 °C until assayed.

All plasma samples were assayed for CNO and clozapine by Precera Bioscience (Franklin, TN) using previously published techniques (Raper et al., 2017). Briefly, liquid chromatography−tandem mass spectroscopy (LC-MS/MS) analyses were performed via reverse phase chromatography using a Shimadzu Nexera X2 UPLC (Columbia, MD) coupled with a QTrap 5500 (Sciex, Framinghman, MA, USA). For plasma samples, 20 μL aliquots of each standard, quality control sample, plasma sample, blank, and double blank were transferred to a labeled 96-well plate. Next, 120 μL of 50 ng/mL internal standard (carbamazepine in acetonitrile) was added to standards, quality controls, plasma samples, and blanks, while 120 μL of acetonitrile was added to the double blank, then the plate was centrifuged for 5 min at 3000 rcf. Then, 75 μL of supernatant was transferred into a labeled 96-well plate, and 75 μL of water was added to all samples. Finally, the plate was heat-sealed for analysis. The calibration range was 1–5000 ng/mL for CNO and 1–1000 ng/mL for clozapine.

### Histology and Visualization of DREADDs

Perfusion and initial preparation of tissue: At the end of the behavioral experiments, the animals received an overdose of pentobarbital, and were transcardially perfused with Ringer’s solution, followed by 4% paraformaldehyde and 0.1% glutaraldehyde in phosphate buffer (PB, 0.2 M, pH 7.4). After removing the brain from the skull, we collected vibratome coronal sections (60 μm) in cold phosphate-buffered saline (PBS, 0.01M, pH 7.4), that were stored at −20° C in an anti-freeze solution (30% ethylene glycol/30% glycerol in PB), until further processing.

Staining of Nissl substance was done to identify the localization of the injection tracks. Following previously published methods (Galvan et al., 2019), localization of hM4Di was revealed using antibodies against the hemagglutinin (HA) tag, which is fused to hM4Di. We selected brain sections containing or adjacent to the virus solution injection tracks, pretreated with 1% normal goat serum, 1% bovine serum albumin and 0.3% Triton X-100, and then incubated for 24 hours in solutions containing either HA-tag antibodies (raised in rabbit, clone C29F4, 1:400, cat # 3724, Cell Signaling Technology, Danvers, MA. As controls, some sections were processed in solutions without the primary antibodies. Incubation in the primary antibodies was followed by 2-hour incubation in anti-rabbit secondary biotinylated antibodies (1:200, Vector Laboratories, Burlingame, CA), and then in avidin-biotin-peroxidase complex (ABC) solution (1:200; Vectastain standard kit, Vector) for 90 min. The sections were then placed in 0.025% 3-3’-diaminobenzidine tetrahydrochloride (DAB, Sigma-Aldrich, St. Louis, MO), 0.01M Imidazole (Fisher Scientific, Pittsburgh, PA) and 0.006% H_2_O_2_ for 10 min. The sections were mounted on slides, cover-slipped, and digitized with an Aperio Scanscope CS system (Aperio Technologies, Vista, CA).

To estimate the extent of the brain regions expressing the hM4Di, we used a series of HA-Tag sections (0.5 mm apart) that encompassed the transfected amygdaloid regions. In digital images (2X), we delineated regions with HA-positive cell bodies, and calculated the area using ImageJ (http://imagej.nih.gov/ij/).

### Behavioral Assessments

Human Intruder Paradigm: The human intruder paradigm was chosen because it has been shown to be a robust task for detecting differences in emotional responses of nonhuman primates after adult or neonatal amygdala lesions (Kalin et al., 2004; Machado and Bachevalier, 2008; Raper et al., 2013b; Raper et al., 2013a). Animals’ responses to the human intruder were assessed under CNO prior to transduction, then again under CNO at 1- and 3-months post-transduction, or under vehicle conditions at 2- and 6-months post-transduction. Clozapine activation of the DREADDs was tested at 5-months post-transduction (Figure 1). On test days when CNO was administered, behavioral testing began approximately 80-90 minutes after injection based on the peak of CNO in cerebrospinal fluid in adult rhesus monkeys (Raper et al., 2017). On test days when clozapine was administered, testing began approximately 5-10 minutes after injection because clozapine readily crosses the blood brain barrier (Baldessarini et al., 1993). During testing, monkeys were separated from their cagemate, transported to a testing room, and transferred to a stainless-steel testing cage (53 × 53 × 55 cm) with one wall made of clear plastic to allow for unobstructed video recording. The human intruder paradigm consisted of three conditions (alone, profile, stare) presented in the same order to all monkeys for a total duration of 30 min. The experimenter wore a rubber mask depicting a different male face at each test, such that both monkeys saw the same novel human intruder at each time point in the study. Other attempts to ensure novelty of the test throughout the six sessions, included the experimenter wearing a different colored lab jacket and either different curtains hung around the room or using a new room to make the environment less familiar. The human intruder test session started with the monkey remaining alone in the cage for 9 min (alone condition) to acclimatize to the environment and obtain a baseline level of behavior. Then the intruder entered the room and sat 2 m from the test cage, presenting his/her profile to the animal for 9 min (profile condition or no eye contact condition in other experiments). The intruder then left the room while the monkey remained in the cage for a 3 min period, then the intruder re-entered the room, and sat 2 m from the cage, making direct eye contact for 9 min (stare condition). Emotional behavior responses to the intruder were assessed using Observer XT 14 software (Noldus Inc., Wageningen, The Netherlands) and a detailed ethogram (Table 1). One experimenter (JR) with a high degree of both intra-rater reliability (Cohen’s κ = 0.98) and inter-rater reliability (Cohen’s κ= 0.86) with other trained experimenters at YNPRC coded all of the videotapes. The experimenter was blind to the animals’ ligand administration while coding the videos.

**Table 1.**
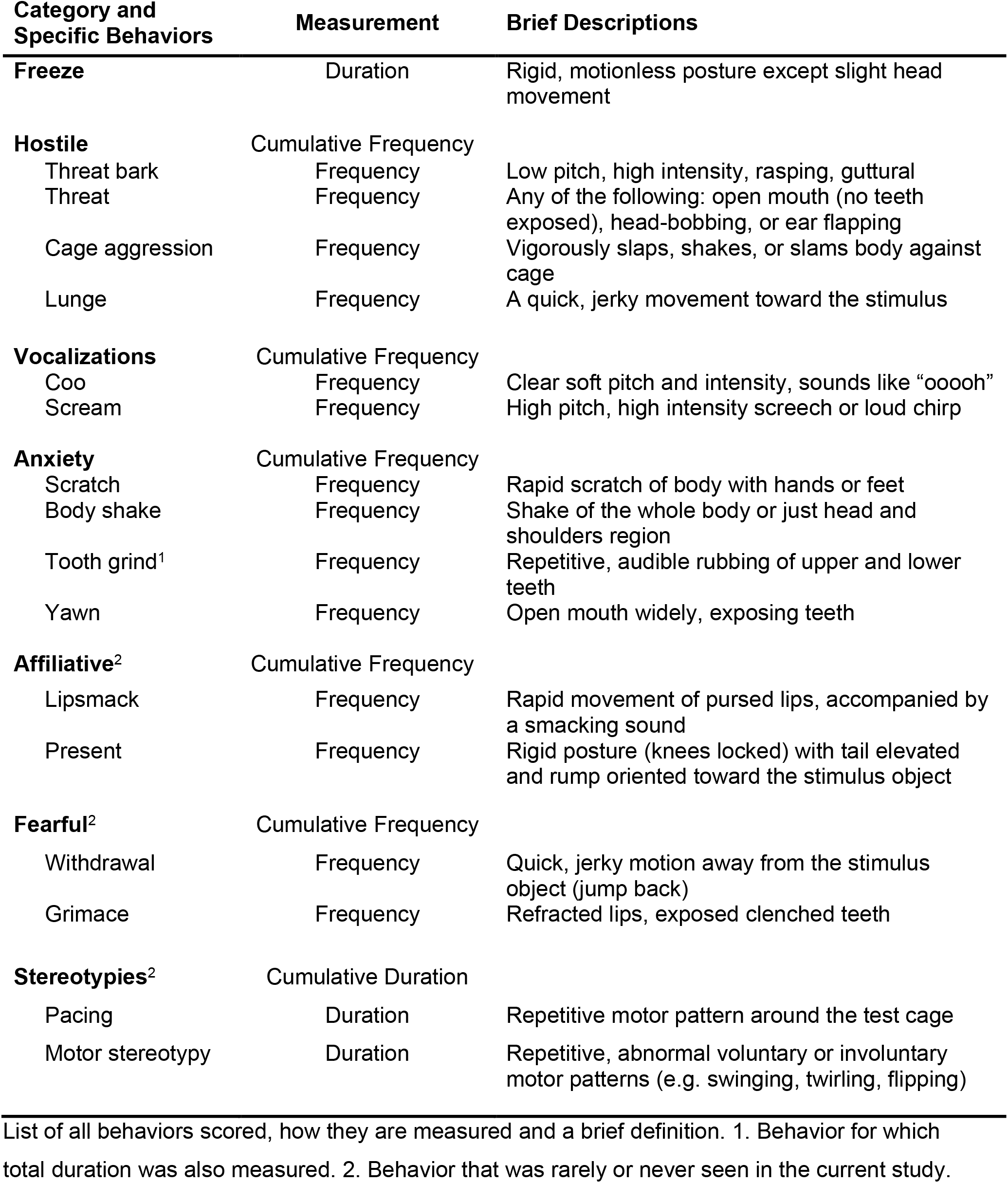
Behavioral Ethogram for Human Intruder

| Category and Specific Behaviors | Measurement | Brief Descriptions |
| --- | --- | --- |
| **Freeze** | Duration | Rigid, motionless posture except slight head movement |
| **Hostile** | Cumulative Frequency |  |
| Threat bark | Frequency | Low pitch, high intensity, rasping, guttural |
| Threat | Frequency | Any of the following: open mouth (no teeth exposed), head-bobbing, or ear flapping |
| Cage aggression | Frequency | Vigorously slaps, shakes, or slams body against cage |
| Lunge | Frequency | A quick, jerky movement toward the stimulus |
| **Vocalizations** | Cumulative Frequency |  |
| Coo | Frequency | Clear soft pitch and intensity, sounds like “ooooh” |
| Scream | Frequency | High pitch, high intensity screech or loud chirp |
| **Anxiety** | Cumulative Frequency |  |
| Scratch | Frequency | Rapid scratch of body with hands or feet |
| Body shake | Frequency | Shake of the whole body or just head and shoulders region |
| Tooth grind<sup>1</sup> | Frequency | Repetitive, audible rubbing of upper and lower teeth |
| Yawn | Frequency | Open mouth widely, exposing teeth |
| **Affiliative<sup>2</sup>** | Cumulative Frequency |  |
| Lipsmack | Frequency | Rapid movement of pursed lips, accompanied by a smacking sound |
| Present | Frequency | Rigid posture (knees locked) with tail elevated and rump oriented toward the stimulus object |
| **Fearful<sup>2</sup>** | Cumulative Frequency |  |
| Withdrawal | Frequency | Quick, jerky motion away from the stimulus object (jump back) |
| Grimace | Frequency | Refracted lips, exposed clenched teeth |
| **Stereotypies<sup>2</sup>** | Cumulative Duration |  |
| Pacing | Duration | Repetitive motor pattern around the test cage |
| Motor stereotypy | Duration | Repetitive, abnormal voluntary or involuntary motor patterns (e.g. swinging, twirling, flipping) |

List of all behaviors scored, how they are measured and a brief definition. 1. Behavior for which total duration was also measured. 2. Behavior that was rarely or never seen in the current study.

Socioemotional Attention Task: Damage to the amygdala has been shown to alter attention toward social cues as well as threats in the environment (Aggleton, 2000; Adolphs et al., 2005; Feinstein et al., 2011; Gothard et al., 2018; Payne and Bachevalier, 2019). To measure changes in attention to social and innate aversive stimuli, subjects’ eye movements were monitored using a Tobii T60/T120 eye tracker (Tobii Technology, Mountain View, CA). Eye tracking technology uses noninvasive infrared light reflections on the subjects’ cornea and retina to measure looking behavior. These reflections were calibrated using a 5-point calibration paradigm at the start of each testing session to control for subtle changes in distance and head position. Subjects were acclimated to an infant nonhuman primate chair (Crist Instrument Co., Hagerstown, MD) and positioned 22” from the eye tracker monitor on which videos were displayed. During test sessions, an experimenter running the eye tracker was visually separated from the monkey by a curtain. The experimenter could monitor whether the tracker was detecting the eyes, and a gaze-trail was displayed over a view of the presented stimuli on the experimenter’s laptop.

Each test session consisted of a novel set of 54 10-second video clips containing Social (36 videos) and Nonsocial (18 videos) content. Videos were presented back-to-back without an intertrial interval between videos clips. All videos were 720 × 480 pixel AVI files, displayed on a 1280 × 1024 resolution monitor (19 × 13 degree of visual angle) with a refresh rate of 120hz. Social videos contained clips of an unfamiliar monkey (4 female, 6 males; 3-20 years age range) looking toward the camera expressing either a threatening, affiliative (lipsmacking), or neutral expression (Mosher et al., 2011; Mosher et al., 2014). Videos were controlled for movement of the camera and luminance (Mosher et al., 2011; Mosher et al., 2014). Nonsocial videos consisted of either aversive stimuli with clips of snakes or spiders, which monkeys innately fear (Mineka et al., 1980; Mineka, 1987; Izquierdo et al., 2005; Chudasama et al., 2009), or neutral stimuli with clips of either enrichment food items (e.g., oranges, grapes, popcorn) or novel items (e.g., train, flower, hot air balloon). Since no head fixation devices were used to restrict the head position to face the monitor, each video was presented twice in each session (to increase the chance that the monkeys would watch the video at least once during the session). Each session lasted between 20-30 minutes. Animals were tested on this socioemotional attention eye tracking task under CNO at 3.5 months, under vehicle conditions at 3.5, 4, and 5.5 months, or under clozapine at 5 months post-transduction (Figure 1). Timing between ligand administration and the beginning of behavioral testing was the same as reported for the human intruder task.

Gaze information was recorded at a rate of 60Hz on a Windows laptop running Tobii Studio software 3.2.2 (Tobii Technology, Mountain View, CA). Several areas of interest (AOIs) were drawn on all video stimuli prior to data collection using the Tobii Studio software. Figure 6A illustrates the AOI for Social videos, regions were defined as eyes (bridge of nose to brow), mouth (nose to chin), face (eyes + mouth), body (outline of monkey body), and background (total viewable movie area). For Aversive and Neutral videos, regions were defined as foreground (outline of object) and background (total viewable movie area). The primary output measure was fixation duration for each pre-drawn AOI. For each video, fixation duration for each AOI was normalized according to the proportion of time spent looking at the corresponding AOI. For example, the average fixation duration in the body AOI was divided by the average fixation duration of the total viewable movie area (background), whereas the mouth or eye AOIs were divided by the average fixation duration of the entire body AOI. Given that animals were not head restrained to face the monitor, most videos were only viewed once out of the two presentations. In cases where the animals looked at the video twice, we used the data from the session with the longest fixation duration. The average proportion of fixation duration was then averaged for each stimulus type (Social: neutral, lipsmack, threat or Nonsocial: neutral, aversive).

### Statistical Analyses

Pharmacokinetic parameters (i.e. area under the curve (AUC)) for CNO and clozapine were determined using Microsoft Office Excel (Microsoft Corporation, Redmond, WA) and results were graphed using GraphPad Prism 7.02 (GraphPad Software Inc., La Jolla, CA). For the human intruder paradigm, first we examined whether CNO prior to transduction impacted behavioral expression on the task by comparing the current data to a group of normally developing controls (n=12; (Raper et al., 2013b)). Using a linear mixed model analysis, we examined the behavioral responses across task Conditions (3: Alone, Profile, Stare) and group (2: CNO prior to transduction in monkeys included in this study, control monkeys from (Raper et al., 2013b) study) as fixed factors and individual monkeys as a random factor. Next, to determine whether CNO or clozapine activation of inhibitory DREADDs could modulate behavioral responses on the human intruder task, we used a linear mixed model analysis examining behavior across task Conditions (3) and ligand (2: CNO/clozapine, Vehicle) as fixed factors and individual monkeys as a random factor. The results obtained with both DREADD ligands (CNO or clozapine) were pooled together for the analysis. Individual behavioral responses across the six human intruder test sessions are shown in Table 2.

**Table 2.** Individual Human Intruder Paradigm Data

| Test Session | Subject | Freezing (seconds) |  |  | Anxiety (frequency) |  |  | Hostility (frequency) |  |  | Vocalizations (frequency) |  |  |
| --- | --- | --- | --- | --- | --- | --- | --- | --- | --- | --- | --- | --- | --- |
|  |  | Alone | Profile | Stare | Alone | Profile | Stare | Alone | Profile | Stare | Alone | Profile | Stare |
| CNO Prior to Transduction | M | 0 | 313.34 | 0 | 0 | 0 | 28 | 0 | 0 | 151 | 0 | 0 | 41 |
|  | F | 0 | 203.97 | 0 | 0 | 0 | 26 | 0 | 0 | 193 | 0 | 0 | 80 |
| CNO - 1 | M | 1.63 | 170.26 | 0 | 0 | 0 | 15 | 0 | 0 | 158 | 0 | 0 | 45 |
|  | F | 0 | 33.83 | 0 | 0 | 0 | 2 | 2 | 0 | 157 | 0 | 0 | 47 |
| Vehicle - 1 | M | 0 | 342.97 | 0 | 0 | 0 | 71 | 0 | 1 | 112 | 0 | 0 | 19 |
|  | F | 0 | 222.58 | 0 | 0 | 0 | 38 | 8 | 0 | 148 | 0 | 0 | 33 |
| CNO - 2 | M | 0 | 72.9 | 0 | 0 | 0 | 33 | 0 | 0 | 87 | 0 | 0 | 16 |
|  | F | 0 | 40.81 | 0 | 0 | 0 | 7 | 4 | 1 | 86 | 0 | 0 | 0 |
| Clozapine | M | 0 | 68.07 | 0 | 0 | 0 | 26 | 0 | 0 | 124 | 3 | 0 | 67 |
|  | F | 0 | 53 | 0 | 0 | 0 | 0 | 1 | 3 | 147 | 0 | 0 | 116 |
| Vehicle -2 | M | 0 | 250.54 | 0 | 0 | 0 | 32 | 8 | 1 | 62 | 8 | 0 | 48 |
|  | F | 0 | 68 | 0 | 0 | 0 | 15 | 2 | 1 | 167 | 39 | 162 | 100 |

List of behaviors scored on the Human Intruder paradigm and the individual values for the male (M) and female (F) infant monkeys across the 6 test sessions.

For the socioemotional attention task, we examined whether CNO or clozapine activation of inhibitory DREADDs could modulate looking patterns across social and nonsocial video stimuli. For social video stimuli, we used a linear mixed model analysis with video Valence (3: Neutral, Lipsmack, Threat) and ligand (2: CNO/clozapine, Vehicle) as fixed factors and individual monkeys as a random factor for each area of interest separately. For nonsocial video stimuli, we used a linear mixed model analysis with video Valence (2: Neutral, Negative) and ligand (2) as fixed factors and individual monkeys as a random factor. Individual responses across the five socioemotional attention test sessions is shown in Table 3. As for the human intruder experiments, the results obtained with CNO or clozapine were pooled together for the analysis. All behavioral data were analyzed using SPSS 24 for Windows (IBM Corporation, Armonk, NY), significance was set at p < 0.05, and effect sizes were calculated using partial eta squared (η_p_^2^).

**Table 3.** Individual Socioemotional Attention Task Data

| Test Session | Subject | Mouth |  |  | Eyes |  |  | Body |  |  | Object |  |
| --- | --- | --- | --- | --- | --- | --- | --- | --- | --- | --- | --- | --- |
|  |  | Neutral | Lipsmack | Threat | Neutral | Lipsmack | Threat | Neutral | Lipsmack | Threat | Neutral | Aversive |
| Vehicle - 1 | M | 8.55 | 7.39 | 1.81 | 4.05 | 16.69 | 25.60 | 26.47 | 23.46 | 28.28 | 28.62 | 56.34 |
|  | F | 0.38 | 4.11 | 1.79 | 22.46 | 24.20 | 29.65 | 48.89 | 28.23 | 44.36 | 68.39 | 60.84 |
| CNO | M | 10.19 | 10.62 | 14.80 | 13.68 | 15.34 | 6.50 | 43.55 | 26.29 | 38.79 | 78.38 | 76.12 |
|  | F | 7.39 | 10.86 | 14.37 | 42.56 | 30.23 | 19.61 | 43.99 | 50.73 | 45.63 | 87.09 | 87.58 |
| Vehicle - 2 | M | 7.85 | 2.82 | 0.59 | 20.00 | 19.41 | 29.39 | 31.27 | 35.83 | 30.92 | 57.21 | 51.49 |
|  | F | 7.81 | 3.44 | 9.02 | 6.88 | 11.06 | 16.05 | 45.54 | 33.40 | 22.54 | 54.46 | 33.69 |
| Clozapine | M | 4.77 | 11.72 | 13.27 | 15.85 | 12.72 | 10.11 | 23.46 | 23.60 | 22.62 | 37.18 | 68.17 |
|  | F | 5.78 | 11.74 | 17.46 | 29.96 | 17.46 | 10.35 | 49.62 | 33.10 | 49.04 | 60.87 | 75.96 |
| Vehicle -3 | M | 14.08 | 2.50 | 4.16 | 16.23 | 14.72 | 9.61 | 21.05 | 32.73 | 43.98 | 59.31 | 44.43 |
|  | F | 11.33 | 10.81 | 11.94 | 26.13 | 19.53 | 15.20 | 50.85 | 56.39 | 62.52 | 44.43 | 29.28 |

Percent looking time at specific areas of interest (AOIs) for Social stimuli containing neutral, lipsmack, or threatening emotional valences as compared to Nonsocial stimuli containing neutral or aversive objects. Individual values for the male (M) and female (F) infant monkeys across the 5 test sessions of the Socioemotional Attention task.

## Results

### Plasma Concentrations of Ligands

Although CNO is the most widely used ligand for activating the DREADDs, it has been shown to have poor penetration of the blood brain barrier and can be metabolized into clozapine (Gomez et al., 2017; Raper et al., 2017; Allen et al., 2018). Importantly, clozapine has a high affinity for the DREADDs and has been shown to preferentially activate these receptors after low dose administration (Armbruster et al., 2007; Gomez et al., 2017). To help interpret DREADD mediated behavioral effects, we collected plasma samples before and after behavioral testing sessions with CNO or low dose clozapine administrations. The two infant monkeys had similar pharmacokinetic profiles for 10mg/kg doses of CNO (Figure 2A). Infants plasma concentrations of CNO were high throughout sample collections with an average AUC value of 171.66 ng/ml·h. As observed previously in adults (Raper et al., 2017), infants also readily metabolized CNO into clozapine, which was detectable starting at 20 minutes post-injection with an average AUC value of 2.03ng/ml·h. Since developmental changes in physiology can impact drug pharmacokinetics and pharmacodynamics (Milsap and Jusko, 1994; Lu and Rosenbaum, 2014), it is important to note that the conversion ratio of CNO to clozapine in infant monkeys (1.1%) was similar to that of adult monkeys (1.3%; (Raper et al., 2017)). Low dose clozapine (0.1mg/kg) administration also exhibited a similar pharmacokinetic profile between the two infant monkeys, however clozapine was not detectable until the second sample collection at 40 minutes post-injection, and plasma concentrations of clozapine were below levels of converted clozapine after 10mg/kg CNO administration (Figure 2B). The time course of clozapine levels in plasma was in agreement with previous reports (Baldessarini et al., 1993).

**Figure 2.**
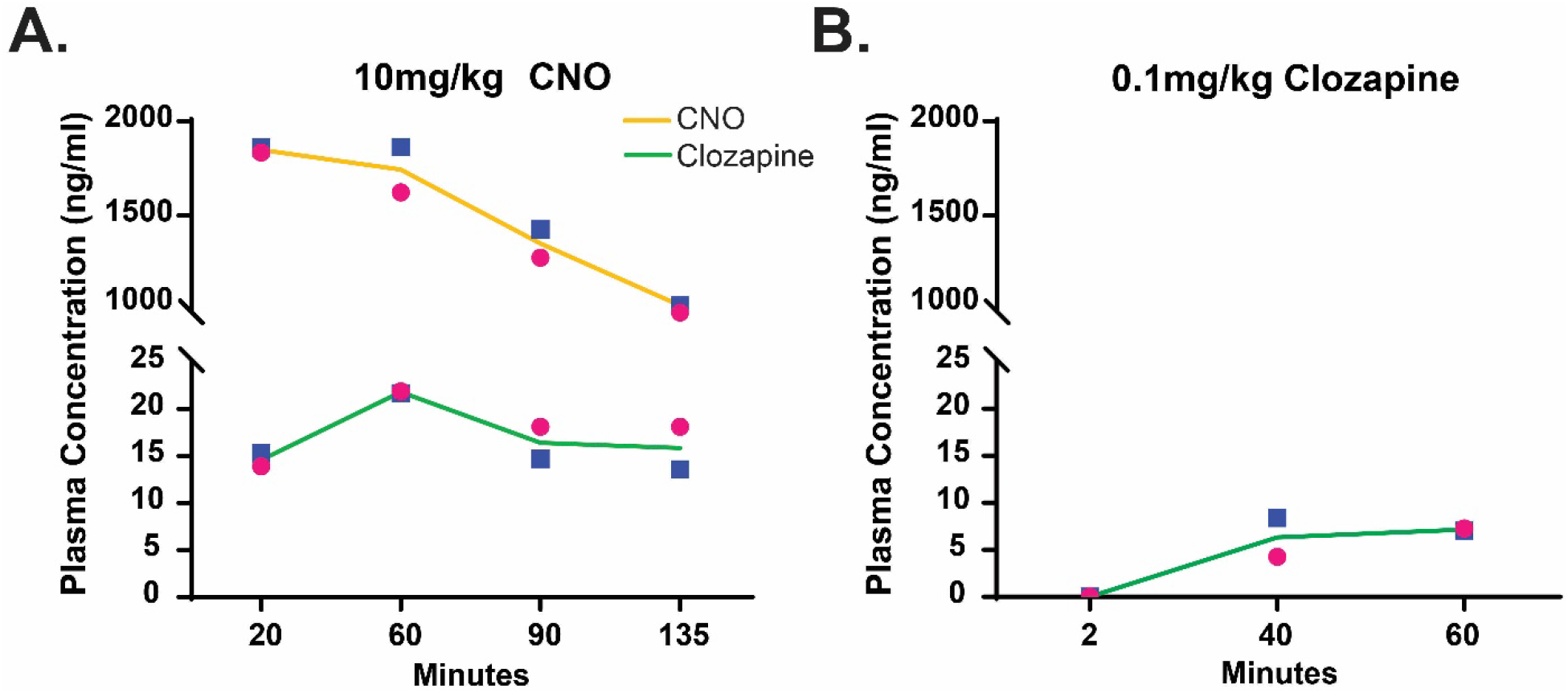
Time-concentration profiles of CNO and clozapine in infant monkeys. Plasma concentrations of CNO (yellow line) and its metabolite clozapine (green line) following SC administration of CNO at 10mg/kg (a) and clozapine at 0.1mg/kg (b) in the infant male (blue square) and female (pink circle) monkeys.

### Structural MRI Scans and Post-mortem Verification of DREADD Expression

While the pre-surgical T1-weighted MRI scan did not reveal any structural abnormalities in the monkeys (Figure 3A), a second scan performed at 4 months post-surgery, showed a reduction in the volume of the left amygdala in the infant female subject (Figure 3B). Since the female’s right amygdala and both hemispheres in the male monkey looked normal in the post-surgical MRI scan, we presume that the reduced volume was due to a unilateral lesion of the amygdala, likely caused by a hemorrhage or other ischemic event associated with the intracerebral injection surgery, rather than as a result of the DREADD expression. The unilateral lesion was confirmed during histology (see inset Figure 3B). Given the extent of the tissue damage in the amygdala region of the left hemisphere, we were unable to identify the injection sites or the DREADD expression in the left hemisphere of the female subject.

**Figure 3.**
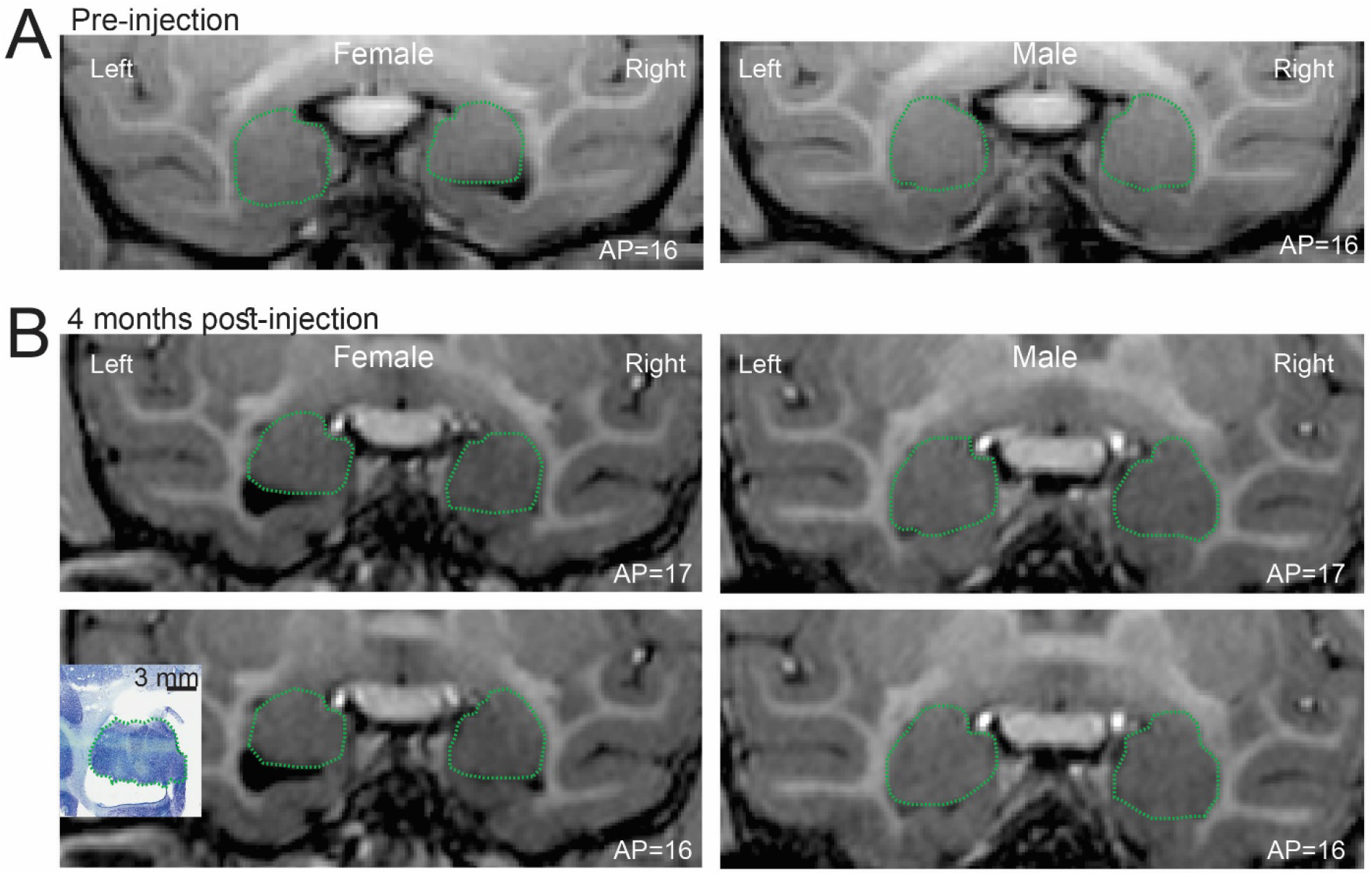
In vivo neuroimaging in infant rhesus monkeys. T1-weighted MRI scan pre-surgery (A) and again at 4 months post-surgery (B) for the female and the male subjects. In the left hemisphere of the female, a reduction in the amygdala was observed at 4 months post-surgery (B), which was later confirmed in the Nissl section (inset). The green dashed lines approximately outline the amygdala.

In Nissl stained sections, we verified that the virus injection tracks were localized in the amygdala (Fig 4A). To identify the areas with robust DREADD expression, we used antibodies against the HA-tag and revealed these antibodies using biotinylated antibodies and immunoperoxidase. In the three hemispheres examined, the HA labeling was found in the basal, lateral, and anterior basal compartments of the amygdala (grey ovals in Fig. 4A). The HA-positive area extended approximately 4 × 3 × 3 mm in the antero-posterior, dorso-ventral and latero-medial planes, respectively. Figure 4B-D show representative images of brain sections close to the viral vector injection sites in the three hemispheres examined. The HA labeling was observed in numerous cell bodies and proximal dendrites of neurons.

**Figure 4.**
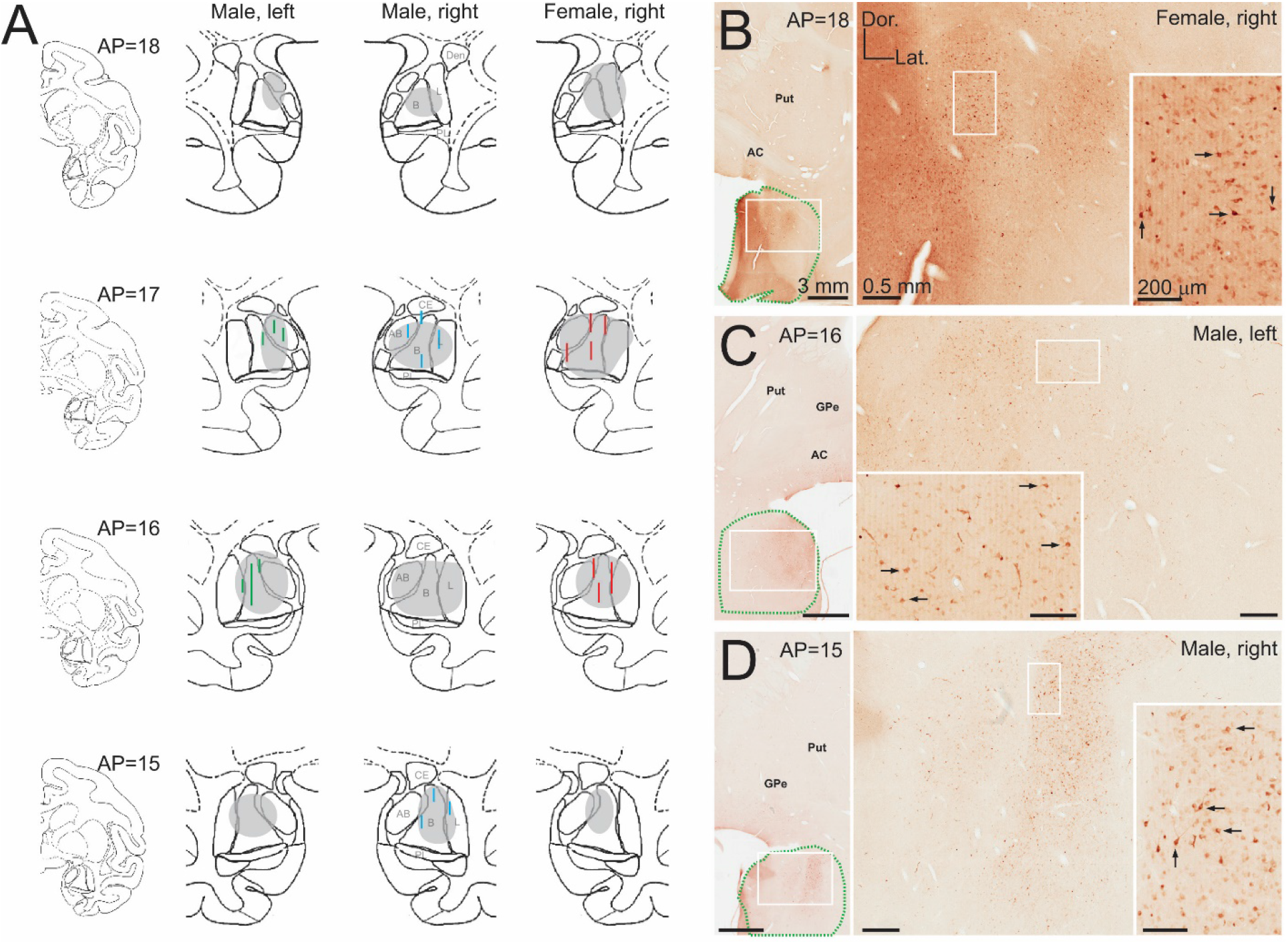
Expression of DREADDs after virus injections in the amygdala. A) Injection tracks (blue or red lines) identified in Nissl stains; and DREADD expression coverage (grey outline) in the amygdala, as identified with immunoperoxidase. B-D) Examples of HA immunoperoxidase labeling in the amygdala. White rectangles indicate areas shown at higher magnification in subsequent panels. Arrow point to examples of HA(hM4Di)-positive neuronal cell bodies. In B to D, the approximate outline of the amygdala is indicated by green dashed lines. Scale bars and orientation in B apply to C and D. AP indicates the approximate antero-posterior plane from the interaural line, according to coronal drawings from a normalized infant rhesus monkey brain (J. Bachevalier, unpublished atlas). Abbreviations: AB, anterior basal amygdala; AC, anterior commissure; B, basal amygdala; CE, central amygdala; Dor., dorsal; GPe, external segment of the globus pallidus: L, lateral amygdala; Lat., lateral; PL, paralaminar amygdala, Put, putamen.

### Modulation of Emotional Reactivity

To investigate whether CNO administration had effects on the behavior of the animals in the human intruder task prior to DREADD transduction, their behavior after CNO administration was compared to that exhibited by monkeys of a similar age in a previous study (Raper et al., 2013b). Both groups exhibited increased freezing in the profile condition (Condition: F[2,24] = 15.31, p < 0.001, η_p_^2^ = 0.56; Figure 5A), increased hostility (Condition: F[2,24] = 22.00, p < 0.001, η_p_^2^ = 0.65) and increased anxiety in the stare condition (Condition: F[2,24] = 10.63, p < 0.001, η_p_^2^ = 0.47; Figure 5B), with no interactions or main Group effects (Freezing: F[1,12] = 0.05, p = 0.83, η_p_^2^ = 0.004; Hostile: F[1,12] = 0.13, p = 0.73, η_p_^2^ = 0.01; Anxiety: F[1,12] = 0.55, p = 0.47, η_p_^2^ = 0.04). Once DREADD transduction was completed, inhibition of the amygdala with either CNO or clozapine administration lead to decreased freezing during the profile condition of the human intruder task as compared to vehicle administration (Ligand × Condition: F[2,23] = 9.23, p = 0.001, η_p_^2^ = 0.45; Figure 5C). DREADD-mediated amygdala inhibition also decreased anxiety expression during the stare condition (Ligand × Condition: F[2,23] = 5.54, p = 0.011, η_p_^2^ = 0.33; Figure 5D). In contrast, DREADD inhibition of the amygdala did not impact hostile behavior expression; which was primarily exhibited during the stare condition in both vehicle and CNO/clozapine conditions (Ligand: F[1,23] = 0.001, p = 0.97, η_p_^2^ < 0.001; Condition: F[2,23] = 100.48, p < 0.001, η_p_^2^ = 0.89; Figure 5E). Also, DREADD-mediated amygdala inhibition did not affect vocalizations, such that both animals emitted more vocalizations during the stare condition regardless of vehicle or CNO/clozapine (Ligand: F[1,23] = 1.74, p = 0.20, η_p_^2^ = 0.07; Condition: F[2,23] = 3.56, p = 0.045, η_p_^2^ = 0.24; Figure 5F).

**Figure 5.**
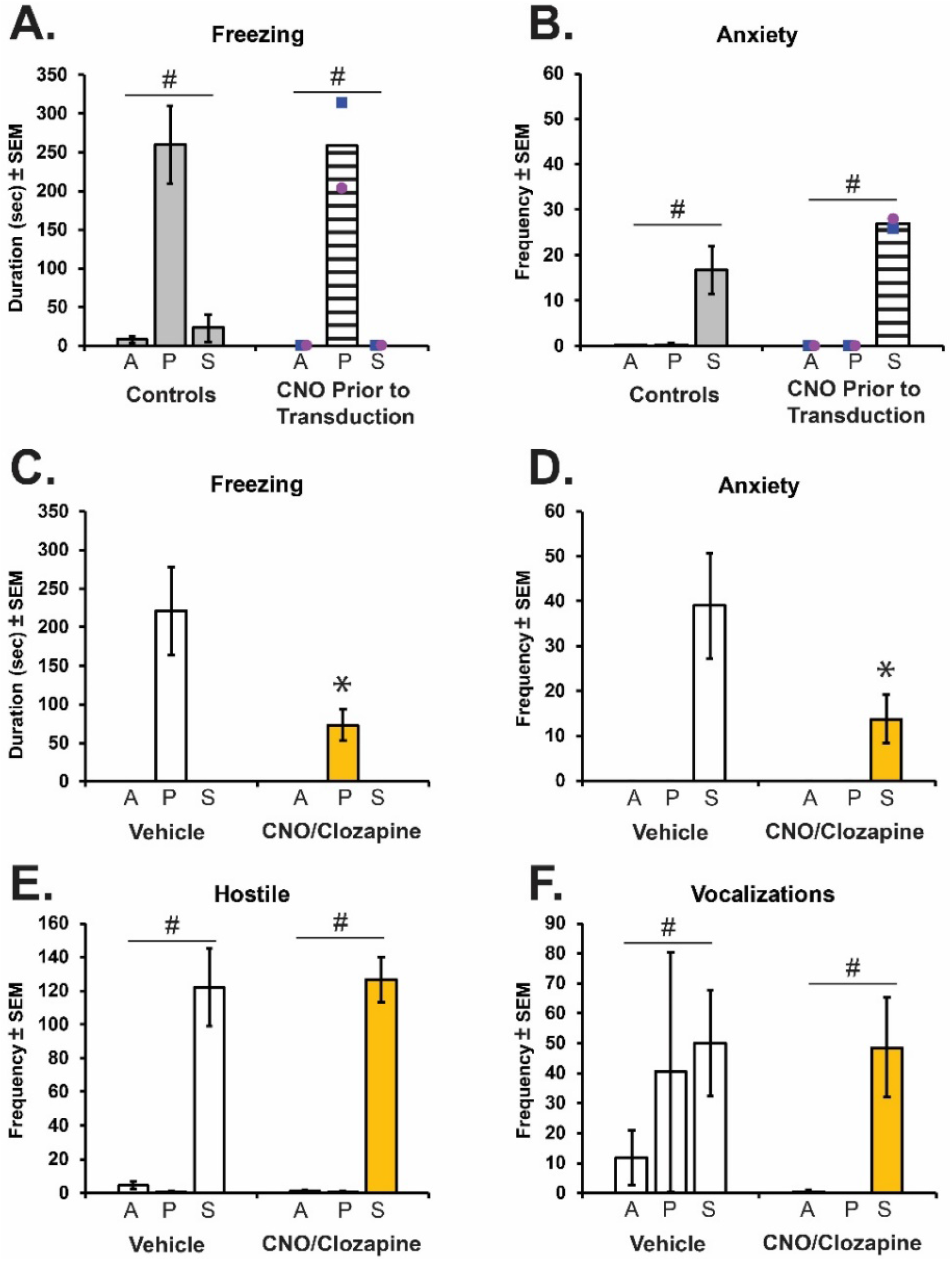
Behavioral responses on the Human Intruder paradigm with and without DREADD inhibition of the amygdala. Freezing (A) and anxiety (B) behaviors measured across task conditions (Alone=A; Profile=P; Stare=S) in the infant male (blue square) and female (pink circle) monkeys prior to transduction (striped bar), as compared to control monkeys from a previous study (“historic controls” = grey bars; (Raper et al., 2013b)). C-F show responses after transduction of the inhibitory DREADDS in the amygdala. Freezing (C), anxiety (D), hostility (E), and vocalizations (F) after CNO or clozapine (CNO/Clozapine, yellow bars) or vehicle (Veh, open bars) administration. # indicates a significant effect of condition on the task, and * indicates a significant Ligand × Condition interaction (p < 0.05).

### Modulation of Socioemotional Attention

The two infants in this study exhibited gaze fixation durations similar to those found in other studies of non-head-restrained infant monkeys (Parr et al., 2016; Ryan et al., 2019). Compared to vehicle sessions, DREADD activation with either CNO or clozapine administration lead to increased attention toward the mouth of conspecifics, depending on the emotional valence of the social video stimuli (Figure 6A). DREADD-mediated inhibition of the amygdala increased attention to the mouth during lipsmack and threatening emotional video clips, but not neutral social video clips (Ligand × Valence interaction: F[2,24] = 3.92, p = 0.034, η_p_^2^ = 0.25; Figure 6B). In contrast, DREADD-mediated inhibition of the amygdala did not impact the amount of time animals spent looking toward the eyes (Figure 6C) or body of conspecifics regardless of the emotional valence (Ligand: F[1,23] = 0.12, p = 0.73, η_p_^2^ = 0.01; F[1,23] = 0.02, p = 0.89, η_p_^2^ = 0.002; Valence: F[2,23] = 0.94, p = 0.40, η_p_^2^ = 0.08; F[2,23] = 0.69, p = 0.51, η_p_^2^ = 0.06, for eyes and body respectively). For nonsocial videos, amygdala inhibition increased attention toward objects (Ligand: F[1,16] = 10.63, p = 0.005, η_p_^2^ = 0.40; Figure 6D) regardless of the stimulus type (Valence: F[1,16] = 0.06, p = 0.81, η_p_^2^ = 0.004).

**Figure 6.**
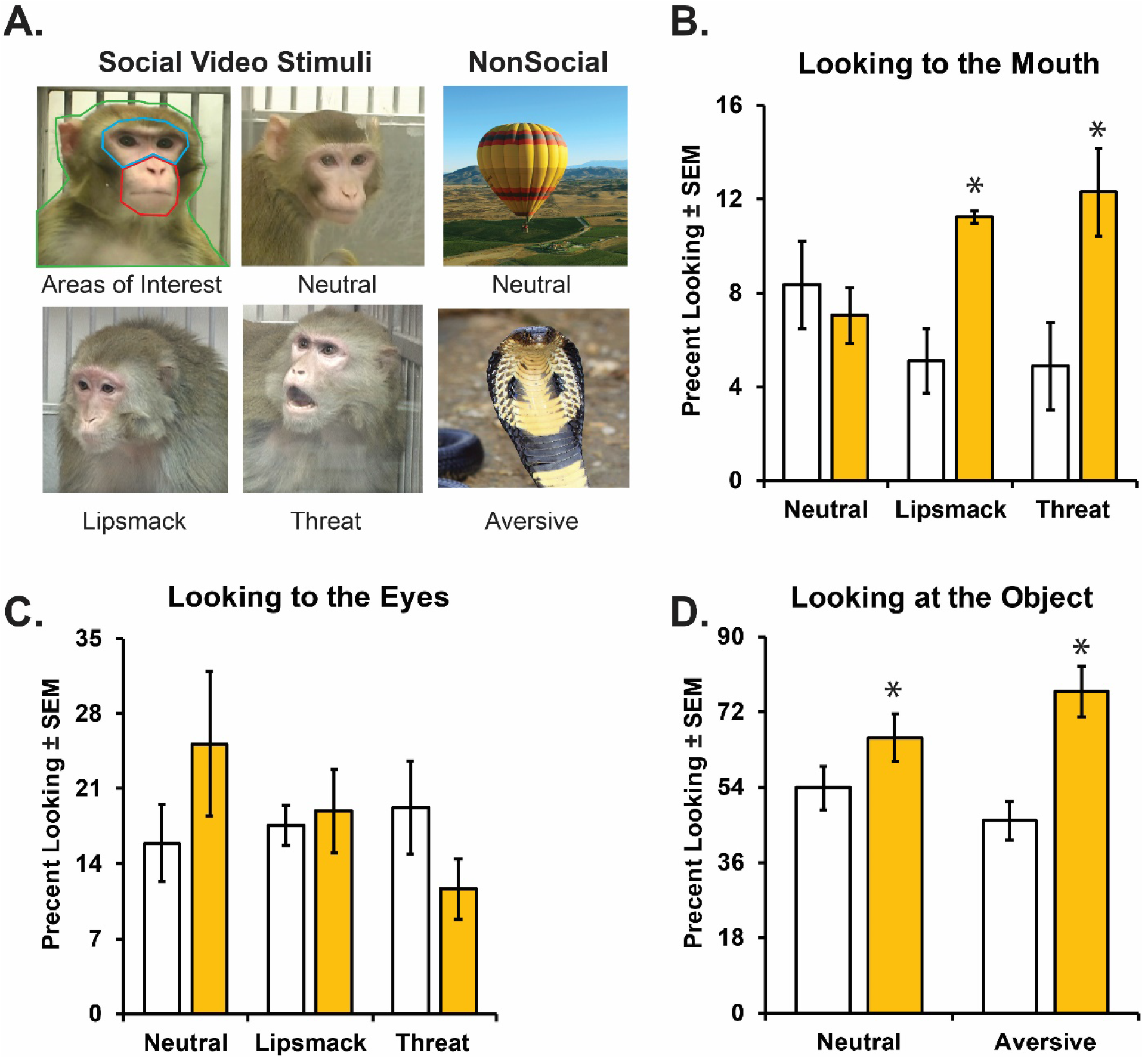
Looking behavior during the Socioemotional Attention task during DREADD-mediated inhibition of the amygdala. Panel A represents examples of the video stimuli type, as well as outlines the specific areas of interests (AOIs) mouth (red), eyes (blue), and body (green) for social stimuli. Percent looking to the mouth (B) or looking to the eyes (C) of Social videos and looking at the nonsocial object (C) after CNO or clozapine (yellow bars) or vehicle (open bars) administration. * indicates a significant difference from vehicle (p < 0.05).

## Discussion

This study provides proof-of-principle that chemogenetic tools can be used in infant nonhuman primates to modulate neuronal activity resulting in transient and reversible changes in behavior. To date nonhuman primate chemogenetic studies have focused on adults (Eldridge et al., 2016; Grayson et al., 2016; Nagai et al., 2016; Upright et al., 2018). Considering its reversibility and low invasiveness, this technique holds great promise for developmental studies in which more invasive techniques cannot be employed. Our study incorporated the use of two amygdala dependent behavioral tasks and investigations of the ability of both CNO and low dose clozapine to activate the DREADD receptors. We have demonstrated that amygdala neurons in infant monkeys can be transduced to express DREADDs after virus solution injections, that the expression is stable for at least 7 months, and that activation of DREADDs in the neurons modulates some anxiety-related behaviors. We have also shown that the reductive metabolism of CNO to clozapine in infant monkeys is similar to adult monkeys, and that low doses of clozapine may be sufficient to elicit the DREADD-mediated behavioral effects. In spite of its technical limitations (see below), our study provides initial data demonstrating that chemogenetics can be used to investigate developmental behavioral neuroscience research questions.

For this study, we chose two tasks that exhibit robust effects after amygdala damage, the human intruder paradigm (Kalin et al., 2004; Machado and Bachevalier, 2008; Raper et al., 2013b; Raper et al., 2013a) and a socioemotional attention task (Adolphs et al., 1994; Adolphs et al., 2005; Chudasama et al., 2009; Machado et al., 2009; Machado et al., 2011; Hadj-Bouziane et al., 2012; Gothard et al., 2018). Given that CNO can be reductively metabolized into the psychoactive compound clozapine (Gomez et al., 2017; Raper et al., 2017; Allen et al., 2018), it was important to establish that CNO administration alone would not impact behavior prior to introducing DREADDs into the amygdala. We confirmed that prior to DREADD transduction in the amygdala, the responses on the human intruder task after CNO administration were similar to those shown in normally developing controls (Figure 5A-B; (Raper et al., 2013b)). Activation of hM4Di DREADDs in the amygdala results in behavioral alterations similar to those seen after both permanent lesion or transient inactivation (Kalin et al., 2001; Kalin et al., 2004; Machado and Bachevalier, 2008; Chudasama et al., 2009; Machado et al., 2009; Bliss-Moreau et al., 2011b; Raper et al., 2013b; Raper et al., 2013a; Dal Monte et al., 2015; Wellman et al., 2016). After the viral DREADD transduction, CNO or low dose clozapine administration reliably decreased freezing and anxiety on the human intruder task. Considering the differences in plasma clozapine levels after CNO or low dose clozapine administration, the current behavioral effects support the finding that very low doses of clozapine are highly effective for DREADD activation (Gomez et al., 2017). Decreased freezing and anxiety in the human intruder task have been reported following amygdala lesions in nonhuman primates during both adulthood (Kalin et al., 2004; Machado and Bachevalier, 2008) or infancy (Raper et al., 2013b; Raper et al., 2013a). In contrast to results obtained after neonatal amygdala lesions on the human intruder task (Raper et al., 2013b; Raper et al., 2013a), the present study in infant monkeys did not exhibit changes in vocalizations and hostile behavior expression after DREADD-mediated amygdala inactivation. The differences between the previous lesion studies and the current study suggest that the more extensive behavioral changes with permanent lesions could be due to lesion-induced plasticity changes after early permanent damage (Machado et al., 2008; Raper et al., 2014b; Grayson et al., 2017; Payne et al., 2017).

Amygdala damage has been shown to alter social behavior and attention to social cues (Adolphs et al., 1994; Bauman et al., 2004; Spezio et al., 2007; Raper et al., 2014a; Dal Monte et al., 2015; Wellman et al., 2016; Bliss-Moreau et al., 2017). Lesions of the amygdala have also been shown to decrease fear reactivity and increase visual interest toward aversive stimuli, such as snakes (Kalin et al., 2004; Chudasama et al., 2009; Machado et al., 2009; Bliss-Moreau et al., 2010; Bliss-Moreau et al., 2011a; Bliss-Moreau et al., 2011b; Feinstein et al., 2011). Using an eye tracking task, we examined how DREADD inhibition of the amygdala altered attention toward social and aversive stimuli. We found that amygdala inhibition with either CNO or clozapine administration resulted in increased looking toward the mouth of novel conspecifics in social stimulus videos. Our findings of increased looking toward the mouth of social videos is similar to results obtained after amygdala damage in adult humans and nonhuman primates (Adolphs et al., 1994; Spezio et al., 2007; Hadj-Bouziane et al., 2012; Dal Monte et al., 2015). A recent study found that adult monkeys with neonatal amygdala lesions looked equally to the mouth and eyes of a conspecfic, whereas controls attended more to the eyes (Payne and Bachevalier, 2019). Those results are similar to the current study, demonstrating that amygdala inhibition only increased the percent looking toward the mouth and did not alter looking at the eyes or body of conspecifics. In addition, DREADD-mediated amygdala inhibition resulted in increased looking at nonsocial videos, including aversive videos containing snakes and spiders. Previous amygdala lesion studies have shown increased interest in aversive stimuli, including increased visual investigation, approach, and manipulation objects (Kalin et al., 2004; Izquierdo et al., 2005; Chudasama et al., 2009; Machado et al., 2009; Bliss-Moreau et al., 2010; Bliss-Moreau et al., 2011a; Bliss-Moreau et al., 2011b; Feinstein et al., 2011). The similarity between the current results and previous lesion research suggest successful chemogenetic inhibition of the amygdala in infant monkeys. In contrast to the differences between DREADD-mediated amygdala inhibition and permanent lesions in response to the human intruder, results from the socioemotional attention task are quite similar to that of permanent lesions (Chudasama et al., 2009; Hadj-Bouziane et al., 2012; Dal Monte et al., 2015). This difference suggests that socioemotional attention is more easily modulated by transient inactivation of the amygdala, whereas behavioral responses, such as hostility, are less sensitive to the temporary DREADD inhibition, as observed during the human intruder task.

There are limitations to this study. First, we only used two infant monkeys. However, this limitation was mitigated by the within-subjects design of the study in which the same behaviors were measured several times under vehicle or CNO/clozapine administration. Second, we only conducted a pre-surgery test of CNO on the human intruder task and not the socioemotional attention eye tracking task. Although, it is possible that CNO/clozapine could have impacted social attention, this is unlikely considering a recent study demonstrating that low dose clozapine does not impact social behavior or working memory in naïve rats (Ilg et al., 2018). Future studies should include testing the DREADD actuators in naïve animals during relevant behavioral tasks. Furthermore, although clozapine at the low doses used here preferentially activated DREADDs, we cannot completely rule out contributions of other neurotransmitter receptors, therefore, future studies should investigate the use of novel inert ligands with high affinity for DREADDs (Bonaventura et al., 2018).Third, there was an unintended amygdala lesion in the infant female subject. Although we cannot rule out the possibility that the lesion was caused by the virus injection or the introduction of the artificial DREADD receptors, this is unlikely given the damage occurred only in one subject and was localized in one hemisphere. Instead, it was most likely due to a hemorrhage or other ischemic event that occurred as a consequence of the intracerebral injections. This event highlights the importance of conducting a post-surgical MRI. Future studies should consider doing a T2-weighted Fluid Attenuated Inverse Recovery (FLAIR) MRI sequence within 5-10 days after surgery which could help to detect edema from a hemorrhage or ischemic event (Nemanic et al., 2002). The extent to which the unilateral lesion impacted behavioral results is unclear; however previous studies have shown that unilateral amygdala lesioned monkeys behave similarly to controls (Kalin et al., 2004; Raper et al., 2013b). Therefore, regardless of whether the bilateral silencing of the amygdala was mediated by DREADDs (infant male subject) or achieved by a combination of unilateral lesion and DREADD activation (infant female subject), the transient and reversible effects on behavior were similar between both subjects (see Figures 5 and 6; Tables 2 and 3). Despite its limitations, the current study is the first to demonstrate that chemogenetic tools can be used in young infant nonhuman primates to address developmental behavioral neuroscience questions.

In summary, transient and reversible inhibition of the amygdala is possible in infant monkeys using chemogenetics, specifically hM4Di DREADDs. This study provides initial data demonstrating that chemogenetics can be used in young animals to investigate developmental neuroscience questions that could not previously be investigated due to the limitations of permanent lesions or invasiveness of pharmacological inactivation studies.

## Acknowledgements

Authors would like to thank Dr. Jocelyne Bachevalier for assistance with viral vector surgery, as well as Jean-Francois Pare and Susan Jenkins for expert technical help with histology. Thanks to Dr. Katalin Gothard for the use of her conspecific video stimuli, as well as Shanice Wilson, B.S., Madi Seaver, Noa Shapiro-Fanklin, B.S., and Sahrudh Dharanedra, B.S. for drawing areas of interests on eye tracking stimulus videos. Many thanks to Drs. Ami Klin and Warren Jones for supporting this research. Thanks to Dr. Thomas Vanderford and Oliva Delmas, B.S. at the Yerkes Virology Core for conducting the neutralizing antibody assay. Thanks to Dr. Boris Kantor at the Duke University Viral Vector Core for the plasmids containing the DREADD construct.

## Notes

**Conflicts of Interest**: None, authors have no conflicts of interest to report.

**Funding Sources**: National Institute of Mental Health (P50-MH081756). Yerkes National Primate Research Center is supported by NIH/Office of the Director P51-OD011132.

#### Summary of Updates

New statistical analyses, Figures, and Tables

